# Mapping the Regulatory Architecture of Circadian Clock Adaptation: A Genome-Wide eQTL Analysis in *Drosophila melanogaster*

**DOI:** 10.1101/2025.05.18.654706

**Authors:** Maya Yair, Bettina Fishman, Maya Aslan, Eran Tauber

## Abstract

The circadian clock system enables organisms to synchronize internal daily rhythms with environmental cues, critically impacting survival and fitness. While the molecular architecture of this system in *Drosophila melanogaster* is well-characterized through transcription-translation negative feedback loops involving ten core clock genes, regulatory genetic variants affecting their expression remain largely unexplored. This study leveraged natural variation in clock gene expression to identify expression quantitative trait loci (eQTLs) through genome-wide association mapping. We utilized the Drosophila Genomic Reference Panel (DGRP), consisting of 205 fully sequenced inbred lines, and measured relative expression levels of all core clock genes in 120 lines via qPCR at a single time point (two hours after light onset). GWAS analysis identified 137 significant SNPs (p < 10^−5^) associated with expression variation across the clock genes. Expression levels showed substantial natural variation, with *pdp1ε* exhibiting the highest variability (89-fold difference between extreme lines) and *cyc* the lowest (11.3-fold). Interestingly, only three significant SNPs were located within clock genes themselves (all in *Clk*), while the majority represented trans-eQTLs in genes with diverse molecular functions. Notable candidates include transcription factors (e.g. *Abd-B, tai, E5*), RNA-binding proteins (PUM, Bru-3, Mbl), and long non-coding and antisense RNAs. Variants were also found in *Mad* and *gbb*, genes implicated in the BMP signaling pathway, which has previously been linked to circadian regulation. This comprehensive eQTL map provides new insights into the regulatory architecture of the circadian clock system and potential mechanisms of evolutionary adaptation to environmental timing cues.

## 1 Introduction

The capacity of organisms to align their daily physiological and behavioral rhythms with environmental cycles represents one of the most remarkable examples of evolutionary adaptation. This temporal synchronization, driven by the circadian clock system, enables anticipation of predictable environmental changes, thereby optimizing survival and reproductive success across diverse ecological contexts (Helfrich-Förster et al. 2020). The molecular architecture underlying these rhythms has been extensively characterized in *Drosophila melanogaster*, revealing a complex network of interlocked transcription-translation feedback loops (TTFLs)(Dubowy and Sehgal 2017). However, the mechanisms governing natural variation in clock gene expression and their evolutionary significance remain incompletely understood.

At the core of the *Drosophila* circadian system lies a conserved transcriptional-regulatory network organized into interlocked feedback loops. The primary loop involves the transcriptional activators CLOCK (CLK) and CYCLE (CYC), which heterodimerize to drive E-box-mediated expression of core genes, including *period* (*per*), *timeless* (*tim*), *vrille* (*vri*), and *PAR-domain protein 1ε* (*Pdp1ε*). PER and TIM proteins accumulate in the cytoplasm, where phosphorylation by DOUBLETIME (DBT) and CASEIN KINASE 2 (CK2) regulates their stability and nuclear entry. Upon translocation to the nucleus, PER/TIM complexes repress CLK/CYC activity, completing the ∼24-hour cycle. Secondary loops involve the repressor VRI and activator PDP1ε, which fine-tune *Clk* expression through competitive binding to its promoter. A third regulatory layer incorporates CLOCKWORK ORANGE (CWO), a CLK/CYC-induced repressor that competes with PER/TIM for E-box binding, ensuring precise transcriptional phasing. Post-translational modifications—particularly rhythmic phosphorylation and Slimb-dependent ubiquitination—introduce critical delays that stabilize oscillations (Ko et al. 2002). This multilayered circuit, operating in ∼150 pacemaker brain neurons, achieves remarkable precision through dynamic protein turnover and subnuclear reorganization of clock genes during repression phases.

Expression quantitative trait loci (eQTLs) have emerged as powerful tools for investigating the genetic basis of gene expression variation and its role in adaptive evolution (Hill et al. 2021). Unlike traditional quantitative trait loci that link genetic variants to phenotypic traits, eQTLs specifically identify genomic regions that influence gene expression levels. This approach has revolutionized our understanding of molecular adaptation by revealing how regulatory polymorphisms shape phenotypic diversity across populations (Ye et al. 2013). In the context of circadian biology, eQTL mapping provides a unique opportunity to uncover regulatory networks that fine-tune clock gene expression in response to local environmental conditions.

The evolutionary significance of circadian clock regulation is exemplified by several well-documented adaptive polymorphisms. For instance, the insertion/deletion polymorphism in the *tim* gene of *Drosophila* results in two allelic forms (*ls-tim* and *s-tim*) with distinct light sensitivities and diapause behaviors, showing clinal distribution patterns across Europe that suggest directional selection (Tauber et al. 2007). Similar adaptive variations have been identified in *cry* and *per* genes, highlighting the importance of clock gene regulation in environmental adaptation (Sawyer et al. 1997; Pegoraro et al. 2022). Beyond coding sequence variations, adaptive evolution also operates on regulatory elements that modulate gene expression. In butterflies, changes in regulatory regions underlie striking differences in wing color patterns involved in mimicry (Nadeau 2016). Similarly, in *Drosophila*, population-specific differences in the expression of the gene *CG9509*, driven by variation in a noncoding enhancer, suggest that selection can act on gene regulation even when the phenotypic effects are less obvious (Glaser-Schmitt et al. 2013).

The circadian clock system represents an ideal model for studying regulatory evolution due to its well-defined molecular architecture and clear adaptive significance. While the core feedback loops are well characterized, the broader regulatory landscape influencing clock gene expression remains incompletely understood. Mapping eQTLs associated with expression variation can uncover novel upstream regulators and pathways involved in the fine-tuning of clock function. These variants may act in *cis* or *trans*, offering insights into both local regulatory control and long-range interactions within the genome that shape circadian transcription.

The *Drosophila* Genetic Reference Panel (DGRP) provides an invaluable resource for investigating natural genetic variation and its phenotypic consequences (MacKay et al. 2012). Comprising 205 inbred lines derived from a natural population in Raleigh, North Carolina, with fully sequenced genomes, this panel enables genome-wide association studies (GWAS) to identify genetic variants associated with quantitative traits, including gene expression levels.

In this study, we exploited natural variation in the expression levels of ten core clock genes across DGRP lines to perform eQTL mapping via GWAS. Our goal was to identify genetic variants associated with clock gene expression and characterize potential regulatory mechanisms contributing to circadian adaptation. By understanding how genetic variation shapes clock gene expression, we aim to gain insights into the molecular basis of temporal adaptation and the evolution of circadian systems.

## 2 Methods

### 2.1 Fly Stocks and Rearing Conditions

The DGRP consists of 205 inbred lines established from wild-mated females collected in Raleigh, North Carolina. These lines were made isogenic through 20 generations of full-sib mating, and their genomes have been fully sequenced, making them valuable resources for population genomics and genome-wide association mapping. The flies were maintained at 18 °C under 12:12-hour light:dark (LD) conditions in standard fly vials. Each vial contains 5 mL of food prepared with 1290 mL distilled water, 122 g cornmeal, 64 g yeast, 120 g molasses, 10 mL propionic acid, and 27 mL anti-fungal agent (Tegosept).

Prior to sample collection, adult flies from each stock were transferred to fresh vials and maintained at 24 °C under 12:12-hour light:dark (LD) cycles for three days. The adults were then removed, and the remaining eggs and larvae were kept under the same conditions until eclosion (approximately 11 days). Following eclosion, approximately 50 3-5 days old flies were collected at Zeitgeber Time 2 (ZT=2) into 15 ml tubes, snap frizzed in liquid nitrogen, and stored at -80 °C until further processing.

### 2.2 RNA Extraction and Processing

The tubes containing approximately 50 flies from each line were transferred to liquid nitrogen for at least 5 minutes. Heads were separated from bodies through vigorous vortexing and collected on ice. RNA was extracted from the heads using the Direct-zol™ RNA MiniPrep (ZYMO RESEARCH) and eluted in 25 μl of nuclease-free water. Genomic DNA contamination was eliminated using RNase-free DNaseI treatment with TURBO™ DNase (Invitrogen). RNA concentration for each sample was determined using a NanoDrop spectrophotometer, and all RNA samples were stored at -80°C.

### 2.3 cDNA Synthesis and Quantitative PCR

First-strand cDNA synthesis was performed using 800 ng of total RNA with the High-Capacity cDNA Reverse Transcription Kit (Applied Biosystems) on a SimpliAmp Thermal Cycler. Each reaction consisted 2 µL RT buffer, 2 µL random primers, 0.8 µL dNTP, 1 µL reverse transcriptase, 1 µL of RNase inhibitor, and nuclease-free water (total of 20 µL). The reactions were incubated at 25°C for 10 min, followed by 37°C for 2 hr and terminated by 85°C 5 min.

Quantitative PCR was conducted using the QuantStudio® 3 Real-Time PCR system to generate CT values for each sample. Each reaction consisted of 5 μl SYBR Green, 0.3-0.5 μl of forward and reverse primers for each clock gene, 4 μl of 1:20 diluted cDNA, and nuclease-free water to a final volume of 10 μl. Denaturation at 95°C for 20 sec, followed by 40 amplification cycles. Each cycle consisted of 1 sec at 95 °C, followed by 20 sec at 60°C. The expression of the housekeeping gene *RpL32* was measured for data normalization. For primer sequences, see Supplementary Table S1.

### 2.4 Standard Curve Generation and Relative Expression Calculation

To assess reaction efficiency and generate standard curves, a pooled cDNA sample was created by combining cDNA from 20 individual samples and diluting it 1:10. Serial dilutions of the pooled sample were prepared and tested in triplicate to determine CT values. Standard curves were automatically generated in QuantStudio software, with slope, efficiency, and Y-intercept calculations. All PCR efficiencies were above 95%. CT values for each sample were applied to the curve equation to calculate relative expression, which was then normalized based on *RpL32* expression. Relative expression levels were transformed, and outlier data points were removed. After removing lines with extreme expression values, 107-117 lines were retained for GWAS analysis

### 2.5 Genome-Wide Association Analysis

SNPs associated with clock gene relative expression levels were identified using the DGRP2 software (MacKay et al. 2012). Following standard practices for DGRP GWAS studies, a p-value cutoff of 10^−5^ was applied. Minor allele frequency (MAF) and effect cutoffs of 0.05 and 0.1, respectively, were used to identify candidate genes for further investigation. We determined the functions of candidate genes using the FlyBase database.

### 2.6 Transcription Factor Binding Site Analysis

To investigate potential regulatory interactions, transcription factors (TFs) identified in our analysis were examined for predicted binding sites near the loci of associated clock genes. Publicly available ChIP-seq datasets from the ReMap 2022 database (Hammal et al. 2022) were used to identify TF binding sites within promoter regions, defined as the 2000 base pairs upstream of the transcription start site (TSS) of each candidate clock gene. In addition, we analyzed these promoter regions using JASPAR (Rauluseviciute et al. 2024) and TFinder (Minniti et al. 2025) to predict potential TF binding motifs. This integrated approach allowed us to evaluate whether the TFs identified in the eQTL analysis are likely to directly bind and regulate the expression of the associated clock genes.

## Results

The relative expression of the different circadian genes showed substantial variability among the DGRP lines (Figure 1). The highest variability was found in *Pdp1ε*, with an 89-fold difference between extreme lines (lines 732 and 744), while *cyc* showed the lowest variability with an 11.3-fold difference between extreme lines (lines 142 and 392).

**Figure 1.**
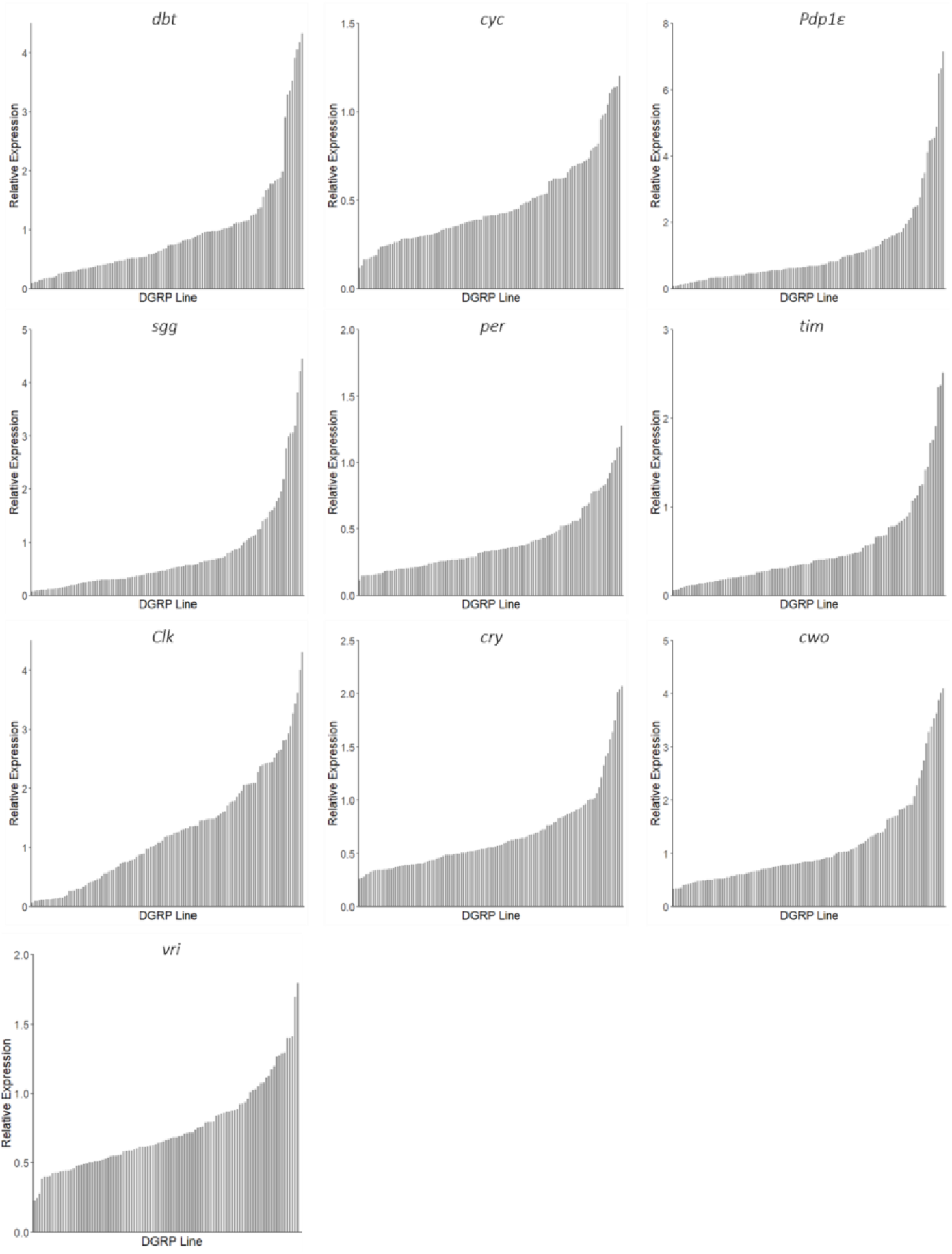
Variation in relative transcript expression of core circadian clock genes across DGRP lines. Relative mRNA expression levels of ten core circadian genes. Expression values were normalized to the housekeeping gene *RpL32* and plotted by DGRP line, ranked by increasing expression for each gene. Each bar represents the normalized expression value for a single inbred line.

GWAS analysis identified 109 significant single nucleotide polymorphisms (SNPs) and 28 indels across the 10 genes examined, using a significance threshold of P = 10^−5^ (Figure 2, Supplementary Tables S2-S3). The majority of identified SNPs were associated with the relative expression of *cry* (34 SNPs) and *per* (23 SNPs). Additionally, 18 SNPs were associated with *Clk* expression, 15 with *cwo*, and 16 with *vri*. Nine SNPs were associated with each of *tim, Pdp1*, and *cyc* expression, while only three and one SNPs were associated with *sgg* and *dbt* expression, respectively.

**Figure 2.**
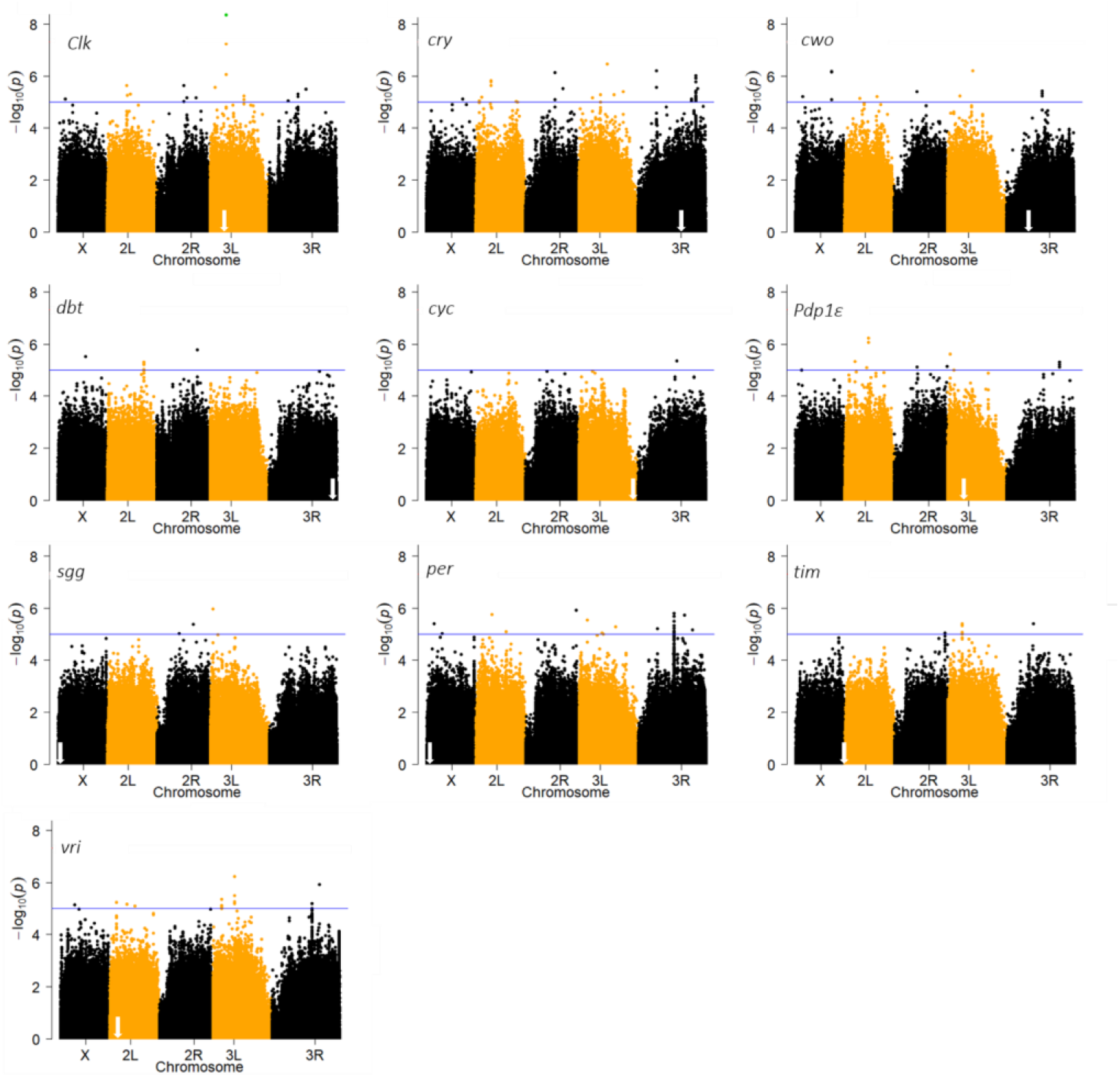
Genome-wide association analysis of clock gene expression. Manhattan plots showing the –log10(p-value) for SNP associations with the relative expression levels of core circadian clock genes across the genome. Each point represents a SNP; its genomic position is plotted along the x-axis, and the strength of association is shown on the y-axis. The blue line marks the nominal significance threshold (p = 1 × 10^−5^). White arrows indicate the approximate chromosomal positions of the tested genes.

Most SNPs were located within intronic regions or upstream and downstream of genes. Eleven SNPs were synonymous substitutions within coding regions of genes such as *hoepel2* (*hoe2*), *CG10073, pumilio* (*pum*), and *CG43394*, which were associated with the relative expression of *Pdp1ε, cyc, per*, and *vri*, respectively. One SNP in the coding region of *Clk* reached genome-wide significance (p = 4.54 × 10^−9^) at the stringent p-value cutoff of 5 × 10^−8^, and another approached this threshold (p = 5.78 × 10^−8^). These two SNPs, along with an additional SNP in the coding region of *Clk*, were the only ones identified within clock genes.

Several genes contained multiple SNPs, including *Abdominal*-*B* (*Abd-B*; 11 variants), *Cytoplasmic linker protein 190* (*CLIP*-190) and *CG32260* (7 variants each), and *sidestep* (side, 6 variants), which were associated with the relative expression of *per, cyc, vri*, and *cry*, respectively.

Additional SNPs were identified upstream of antisense RNAs CR46049, CR4406, and CR44020, as well as in the exon of antisense RNA CR45928, which were associated with relative expression of *per, vri, cry*, and *tim*, respectively. According to FlyBase, the functions of these antisense RNAs remain unknown. Seven SNPs were also identified in long non-coding RNAs (lncRNAs), including lncRNA:CR46006, which was associated with *cry* expression.

Seven significant SNPs were located within introns or in proximal regulatory regions (upstream or downstream) of protein coding genes with currently unknown functions, and an additional 15 SNPs were found in intergenic regions. Strikingly, only a small fraction of the 137 significant variations were mapped within or near the core clock genes themselves. Instead, the vast majority were located in distant genomic regions, often on different chromosomes from the target genes. This spatial distribution strongly indicates that most of the regulatory variants uncovered in our analysis act in *trans*, rather than in *cis*. Many of these trans-eQTLs were located within or adjacent to genes encoding transcription factors, RNA-binding proteins, or lncRNAs, suggesting a broad and distributed regulatory network that influences circadian gene expression beyond the canonical feedback loops.

To visualize the magnitude of genotype-dependent expression differences, we plotted relative expression levels stratified by SNP genotype for four representative variants (Figure 3). The synonymous SNP in *Clk* showed approximately a 2.4-fold difference in median expression between allelic groups. A trans-acting variant in *hoe2* associated with *Pdp1ε* expression showed a 2.1-fold difference. The SNP in *Abd-B*, linked to *per* expression, exhibited a 2-fold effect, while the variant near *Mad* showed a 1.3-fold difference in *vri* expression.

**Figure 3.**
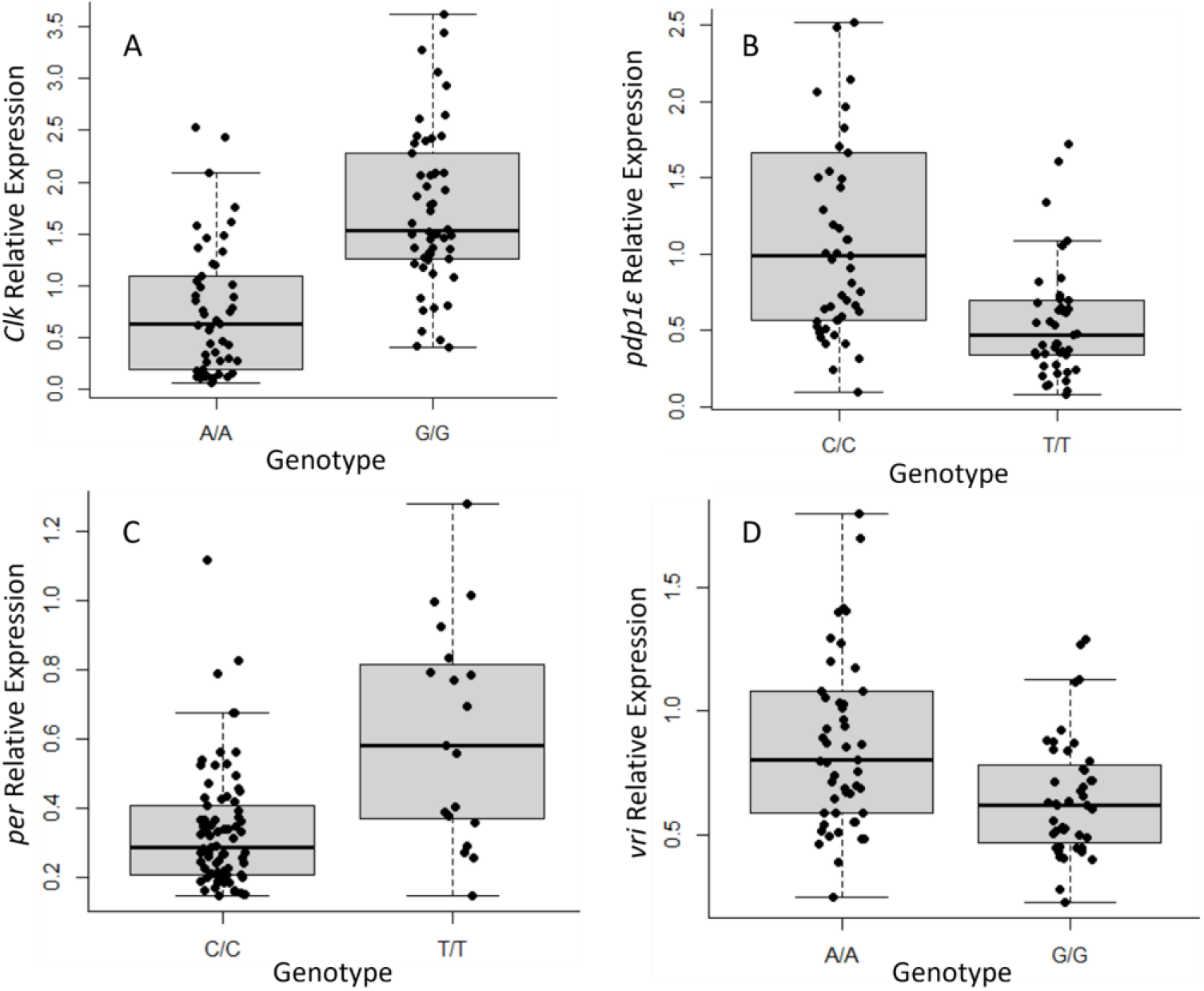
Effect size of selected eQTLs on circadian clock gene expression. Boxplots showing relative expression levels of circadian genes stratified by SNP genotype for four representative variants. (**A**) A synonymous SNP in *Clk* associated with *Clk* expression (cis-eQTL). (**B**) A synonymous SNP in *hoe2* associated with *Pdp1ε* expression. (**C**) A SNP in the homeotic gene *Abd-B* associated with *per* expression. (**D**) A SNP near *Mad*, a transcription factor associated with *vri* expression. Expression values represent normalized transcript levels relative to *RpL32*.

Promoter analysis revealed candidate transcription factor binding sites in several circadian clock genes (Supplementary Table S4). By analyzing previously published ChIP-seq datasets compiled in the ReMap 2022 database, we identified binding sites for multiple transcription factors within the promoter regions of candidate clock genes. Notably, three distinct MAD binding sites were found near the promoters of different *vri* transcript isoforms. These sites were derived from datasets generated in embryonic tissues, embryonic cell lines, and whole adult flies, suggesting potential isoform-specific regulation by MAD. In addition, a single Abd-B binding site was detected within the promoter region of *per*, based on data from whole adult fly samples, indicating possible direct regulation of *per* by Abd-B in adult tissues. The full results of this analysis are summarized in Supplementary Table S4.

Motif analysis using TFinder and the JASPAR database further identified putative transcription factor binding motifs within clock gene promoters (Supplementary Table S5). A single high-scoring Mad motif was located in the *vri* promoter region. Multiple Abd-B motifs were found within the *per* promoter, and several E5 motifs were detected in the *Clk* promoter. All motifs had relative adjusted scores above 0.85, supporting the likelihood of specific and potentially functional TF–DNA interactions (Figure 4).

**Figure 4.**
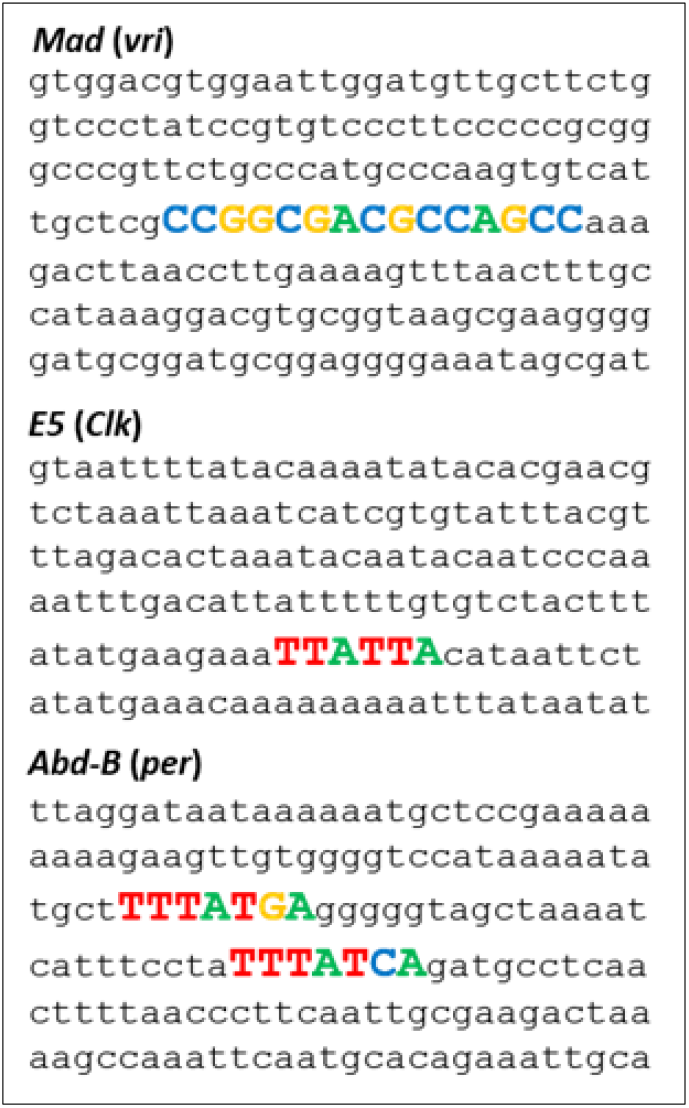
Representative transcription factor binding motifs identified in promoter regions of clock genes. Promoter sequences of circadian clock genes were analyzed to identify potential transcription factor binding sites using JASPAR and TFinder. Only high-confidence sites with relative adjusted scores greater than 0.85 were considered. The figure shows representative predicted binding sites: a Mad site in the *vri* promoter at chromosome 2L:5,288,182; E5 sites in the *Clk* promoter at 3L:7,776,983; and Abd-B sites in the *per* promoter at X:2,684,407 and X:2,684,438. Additional binding sites were identified but are not shown (see main text).

## Discussion

Regulatory variation in gene expression is a key driver of phenotypic diversity and local adaptation. While the core molecular components of the circadian clock have been well characterized in *Drosophila melanogaster*, much less is known about the natural genetic variation that modulates their expression. Here, we used genome-wide association analysis in the *Drosophila* Genetic Reference Panel (DGRP) to map expression quantitative trait loci (eQTLs) influencing the transcription of ten core clock genes. By combining high-resolution genotyping with quantitative expression data from over 100 genetically distinct lines, our study provides a comprehensive view of the regulatory architecture underlying circadian gene expression.

Most identified SNPs were trans-eQTLs, while three cis-eQTLs were detected within the *Clk* locus. This finding was unexpected, as the circadian clock is a complex system where genes and their protein products interact within tightly coupled TTFLs. Similar to well-known regulators of molecular rhythms, the trans-eQTLs were identified in genes with diverse molecular functions, including transcription factors, enzymes, transporters, and structural molecules. These genes play roles in various biological processes, such as cell organization and biogenesis, development, response to stimuli, signaling, transport, and localization.

To narrow down our list of candidate genes, we focused on genes and regulatory elements known to play roles in gene expression. Several notable trans-eQTLs were identified in transcription factors, including *Mothers against dpp (Mad)*, which was associated with the expression of *vri*. MAD is a key component of the highly conserved BMP signaling pathway, which influences synaptic connectivity and regulates transcription. In this pathway, BMP ligands such as *GLASS BOTTOM BOAT (GBB)* bind to type I and type II receptors, forming activated complexes in the presence of dimerized ligands. These complexes initiate intracellular signaling through the phosphorylation and nuclear translocation of MAD and a second transcription factor, MEDEA (MED), which together regulate gene expression. Several studies have shown that the BMP pathway is active in specific circadian pacemaker neurons and modulates neuronal connectivity during the day, thereby influencing the structure and output of the circadian network (Polcowñuk et al. 2021). The reduced behavioral rhythmicity resulting from BMP ligand downregulation, along with lengthening of the circadian period upon overexpression of *Mad* and *Med*, supports a role for BMP signaling in clock regulation. This mechanism is further supported by the identification of MAD and MED binding sites near the *Clk* promoter (Beckwith et al. 2013), and by our finding of three trans-eQTLs associated with *tim* expression in *gbb*, a ligand that activates the BMP pathway. BMP activation may reduce *Clk* expression, thereby decreasing PER levels and slowing the circadian rhythm by increasing the free-running period. An earlier study further supports a regulatory link between the BMP pathway and *vri*, identifying *vri* as a dominant maternal enhancer of *dpp*. Maternal *vri* mutations were shown to exacerbate the embryonic patterning defects and wing abnormalities caused by *dpp* mutations. Moreover, *Mad* mutations enhanced the wing phenotypes of *vri* mutants, suggesting a functional interaction between VRI and MAD in the regulation of developmental processes, including wing formation (George and Terracol 1997).

We also identified an eQTL in *Tamian* (*tai*), a Basic Helix-Loop-Helix (bHLH) transcription factor, which was significantly associated with the relative expression of *Clk*. A recent study demonstrated that *tai* knockdown in both the linden bug (*Pyrrhocoris apterus*) and cockroach (*Blattella germanica*) slowed the locomotor circadian period by 2-5 hours, suggesting a conserved *tai* clock function in certain insect groups (Smykal et al. 2023).

The third transcription factor that harbored an eQTL was *E5*, an NK-like (NKL) homeobox transcription factor (Sundram et al. 2012). This eQTL was associated with the expression of *Clk*. Promoter analysis of *Clk* revealed multiple *E5* DNA binding sites. We also identified eleven SNP in *Abd*-*B*, a transcription factor that is one of the three Hox genes in the bithorax complex and plays a crucial role in regulating the development of posterior abdominal segments, external genitalia, and gonads. The variants in *Abd-B* were associated with *per* transcriptions, and promoter analysis revealed 12 *Abd-B* DNA binding sites.

Our GWAS analysis identified candidate trans-eQTL in *CG2991* associated with *cry. CG2991* encodes an E3 ubiquitin ligase. Several E3 ligases have been previously shown to play roles in the circadian rhythm of *Drosophila* (Srikanta and Cermakian 2021). The E3 ligase SLIMB ubiquitinates PER and TIM, while Circadian Trip (CTRIP) has been identified as increasing the ubiquitination of CLK. In the presence of the E3 ligase JETLAG (JET), CRY and JET form a complex with TIM, preventing nuclear localization and facilitating the ubiquitination of TIM.

Several eQTL were mapped to long non-coding RNA (lncRNA) including lncRNA:iab8 and lncRNA:CR46006, which were associated with the relative expression of *cwo* and *cry*, respectively. lncRNA:iab8 regulates the Hox gene *Abdominal-A* (*abd-A)*, while lncRNA:CR46006 is involved in miRNA-mediated post-transcriptional gene silencing and is a component of the RISC complex. Previous studies have already shown that lncRNAs play a regulatory role in sleep in *Drosophila*, such as lncRNA *yellow-achaete intergenic (yar*) (Soshnev et al. 2011). Therefore, the candidate lncRNAs identified in our analysis could serve as valuable leads for uncovering additional regulatory elements of the circadian system.

Post-transcriptional gene regulation is also mediated by RNA-binding proteins (RBPs), which interact with single-stranded or double-stranded regions of RNA molecules via their binding domain (Gamberi et al. 2006). We identified variants in the genes encoding RNA-binding proteins such as *pumilio* (*pum*), *bruno-3* (*bru*-3), and *Muscleblind* (*Mbl*). Pum is known to exert a repressive effect on the translation of *hunchback* (*hb*), Bru*-3* regulates sarcomeric transcripts, and Mbl modulates RNA metabolism through the regulation of alternative splicing, transcript localization, and the biogenesis of miRNAs and circRNAs. It has been previously demonstrated that RNA-binding proteins such as LARK play a role in the regulation of circadian rhythms (Sundram et al. 2012). LARK is an essential RNA-binding protein required within PDF neurons to maintain robust rhythmicity. Given this, *pum, bru*-3, or *Mbl* may also be involved in the circadian clock, similarly to *lark*.

A previous study that utilized the DGRP to conduct eQTL mapping identified several eQTLs associated with *Clk, sgg*, and *cyc* (Everett et al. 2020). The majority of the identified variants were cis-eQTL. Notably, none of the trans-eQTL was identified in our anlysis. Another study used a different fly panel, the Drosophila Synthetic Population Resource (DSPR) (Smith and Macdonald 2020). Several significant cis- and trans-eQTLs were identified for *cyc, tim, pdp1ε, cwo, sgg*. Notably, none of the eQTLs identified in their analysis overlapped with those identified in our study. On the contrary, in both studies, different variants were identified in *Fibp* that were associated with *cyc* expression. One variant was identified in *Med* and was associated with the expression of *dbt*. The discrepancies between the studies might be due to the different diel times that were used to collect the samples, the panel of flies, and the tissue that was used. Indeed, a recent study showed that substantial variation exists in the expression of clock genes among different tissues (Litovchenko et al. 2021).

The variants identified in our study may represent key starting points in the search for additional regulatory factors of the circadian clock. Follow-up experiments aimed at elucidating the mechanisms through which these genes modulate the clock will provide deeper insights into the well-regulated circadian system at multiple biological levels. Ultimately, determining the functional context of these variants and further investigating their interactions with environmental factors may yield valuable information regarding the adaptive processes they facilitate in the organism, particularly in relation to its local environment.

## Conflict of Interest

None declared.

## Notes

### Competing Interest Statement

The authors have declared no competing interest.

### Funding

- Israel Science Foundation: 2121/23

